# Stabilization of E-cadherin adhesions by COX-2/GSK3β signaling is a targetable pathway in metastatic breast cancer

**DOI:** 10.1101/2022.01.18.476813

**Authors:** Kuppusamy Balamurugan, Saadiya Sehareen, Shikha Sharan, Savitri Krishnamurthy, Wei Tang, Lois McKennett, Veena Padmanaban, Kelli Czarra, Andrew J. Ewald, Naoto T. Ueno, Stefan Ambs, Dipak K. Poria, Esta Sterneck

## Abstract

Metastatic progression and treatment-resistance of breast cancer has been associated with epithelial-mesenchymal-transition including downregulation of E-cadherin (*CDH1*) expression, which can be initiated by inflammatory mediators such as COX-2. Recently, E-cadherin-mediated, cluster-based metastasis and treatment resistance has become more appreciated, though the mechanisms that maintain E-cadherin expression in this context are unknown. Through studies of inflammatory breast cancer and an *in vitro* tumor cell emboli culture paradigm, we identified a role for COX-2, a target gene of C/EBP*δ*, or its metabolite PGE2 in promoting protein stability of E-cadherin, β-catenin and p120 catenin through inhibition of GSK3β, without affecting *CDH1* mRNA. The COX-2 inhibitor celecoxib downregulated E-cadherin complex proteins and caused cell death. Co-expression of E-cadherin and COX-2 was seen in breast cancer patients with poor outcome and, along with inhibitory GSK3β phosphorylation, in patient-derived xenografts of triple negative breast cancer. Celecoxib alone decreased E-cadherin protein expression within xenograft tumors, reduced circulating tumor cells and clusters, and in combination with paclitaxel attenuated or regressed lung metastases. This study uncovered a mechanism by which metastatic breast cancer cells can maintain E-cadherin-mediated cell-cell adhesions and cell survival, suggesting that patients with COX-2+/E-cadherin+ breast cancer may benefit from targeting of the PGE2 signaling pathway.

## INTRODUCTION

Breast cancer (BC) subtypes are classified by expression of hormone receptors (HR) for estrogen and progesterone as well as HER2. Inflammatory breast cancer (IBC) is a rare, highly invasive subtype of BC which can include any of the classical subtypes but does not have IBC-specific treatment options (1). While the term “inflammatory” has been considered a misnomer for IBC, inflammation-associated signaling pathways including NF-κB, COX-2, and JAK/STAT3 signaling are indeed activated in IBC (1-3). These pathways have commonly been associated with induction of epithelial-mesenchymal transition (EMT) of BC cells, which involves downregulation of E-cadherin gene expression to promote invasiveness (4). However, E-cadherin expression is maintained in many advanced breast cancers including IBC, where it plays an important role in the formation of tumor cell emboli within the cancer parenchyma and dermal lymph vasculature and which predict poor outcome (5-7). In addition, it is becoming more evident that cancer cell dissemination may not require complete EMT but rather fluid transitions between EM phenotypes or hybrid states (8, 9, 10). Thus, it has been shown that E-cadherin can contribute to collective cell migration, establishment of metastases, chemotherapy resistance, and cancer cell survival under hypoxia and as circulating tumor cells (CTCs) in clusters (8, 11, 12). Indeed, among BC subtypes, only lobular carcinoma is marked by downregulation of E-cadherin, while most ductal carcinomas maintain high E-cadherin expression (13). This includes metastases (8), which are the main cause of BC mortality, but are still significantly understudied. A detailed understanding of the molecular pathways that maintain E-cadherin expression and cell-cell adhesion in metastatic cells such as during emboli formation will provide new mechanistic insights into BC progression. Experiments with cell culture conditions that were designed to mimic the lymphatic environment showed that IBC but not non-IBC cell lines form emboli-like structures *in vitro*, which resemble emboli in patients (14, 15). This assay system can, thus, be used to interrogate the pathways leading to E-cadherin-mediated cancer cell adhesion (16).

We began this study after observing that the transcription factor C/EBPδ (*CEBPD)* was highly expressed in IBC cell lines and in parenchymal tumor cell emboli of patient tissues. In many cell types, C/EBPδ expression is induced by cytokines via STAT3 and NF-κB signaling and participates in the further induction of pro-inflammatory genes including IL-6 and the IL-6 receptor (17, 18). Within non-IBC breast cancer, high C/EBPδ protein expression is mostly seen in low grade, HR+ luminal-epithelial tumors, and attenuates cell proliferation, motility, and invasion in HR+ cell lines in culture (19). However, in the context of inflammation and hypoxia, C/EBPδ promotes cancer stem cell-associated phenotypes (18). Thus, the role of C/EBPδ depends in part on cell type and context (17). In this report, we show how studies in 3D culture revealed that C/EBPδ supports E-cadherin expression and cell-cell adhesions through expression of COX-2, which sets in motion a signaling cascade that leads to stabilization of epithelial cadherin/catenin proteins. We further provide in vivo evidence that the COX-2/E-cadherin pathway extends beyond IBC, may contribute to poor prognosis in breast cancer, and offers potential for targeted therapy.

## RESULTS

### C/EBPδ is expressed in IBC cells and promotes expression of E-cadherin and cell-cell adhesion in 3D

Because of the implication of inflammation-related signaling pathways in IBC and C/EBPδ’s role in pro-inflammatory signaling (1, 17, 20), we analyzed C/EBPδ expression in IBC tissues by immuno-histochemistry. Analysis of 39 specimens representing different BC subtypes yielded variable C/EBPδ expression patterns and no significant nuclear staining in most tumor cells. However, in 13 of 14 specimens that also contained tumor cell emboli, nuclear C/EBPδ expression was detectable in cells within emboli (**Fig. 1A)**. Our prior analysis of patient-derived xenografts (PDXs) showed that immunohistochemistry with this antibody for C/EBPδ is specific (19) but not very sensitive (18). Thus, while C/EBPδ expression in IBC overall remained unclear, the results indicate that C/EBPδ can be expressed in cells within emboli that have intravasated into the lymphovascular space. Analysis of BC cell lines, however, revealed that C/EBPδ expression was higher in IBC than most of the non-IBC cell lines tested (**Fig. 1B**). In concordance with our previous studies in non-IBC TNBC (18), C/EBP*δ* supported in vitro invasiveness, pro-oncogenic gene expression and cancer cell stemness in SUM149 and IBC-3 cell lines, and growth of established SUM149 experimental metastases in vivo (**Fig. S1A-G**). Because C/EBPδ expression in patient tissues was most pronounced in tumor cell emboli, we next employed a 3D *in vitro* culture model in which cells are seeded in suspension with PEG8000-supplemented media and rocked at slow speed. This paradigm was developed to mimic the mechanophysical environment encountered by the cancer cells within lymphatic vessels (14). Comparison of three IBC cell lines and four non-IBC cell lines showed that under these conditions (from here on referred to as “3D”), only IBC cells aggregate into large, tight clusters (from here on referred to as “emboli”), which resemble tumor cell emboli observed in vivo (14). Compared to adherent cells grown in plastic dishes (2D), culture of SUM149 and IBC-3 cells in 3D induced *CEBPD* mRNA and protein expression despite the significantly different basal levels (**Fig. 1C-D**). In contrast, expression of the related protein C/EBPβ was not induced (**Fig. 1C)**. To test whether C/EBPδ plays any role in the emboli formation, C/EBPδ was silenced in SUM149 and IBC-3 cells prior to 3D culture, which resulted in fewer and/or smaller emboli as significantly fewer cells aggregated in 3D (**Fig. 1E**). Studies have shown that tumor cell emboli depend at least in part on cell-cell adhesions through E-cadherin, which are dependent on binding of Ca^++^ (6). Thus, we assessed the effect of Ca^++^ chelation by EDTA and found that emboli formed by C/EBPδ-depleted cells dissociated more readily when incubated with EDTA (**Fig. 1F-G**). Western analysis revealed that emboli of *CEBPD*-silenced cells contained significantly lower levels of not only E-cadherin protein but also α*−*catenin, β*−*catenin, and p120 (**Fig. 1H)**, which are part of the E-cadherin adhesion complex (6). However, C/EBPδ depletion did not affect the mRNA levels of the corresponding genes (**Fig. 1I**). Next, we asked if this pathway was only necessary for the process of emboli formation or also for the maintenance of established cell-cell adhesions. Doxycycline treatment of established IBC-3 emboli downregulated E-cadherin, β−catenin, and p120 proteins (**Fig. 1J**) and caused partial disintegration of emboli (**Fig. 1K**) in cells with doxycycline-inducible *CEBPD*-targeted shRNA but not in control shRNA expressing cells. Taken together, these data show that C/EBPδ expression in IBC cells supports the expression of E-cadherin complex proteins and cell-cell adhesion.

**Figure 1.**
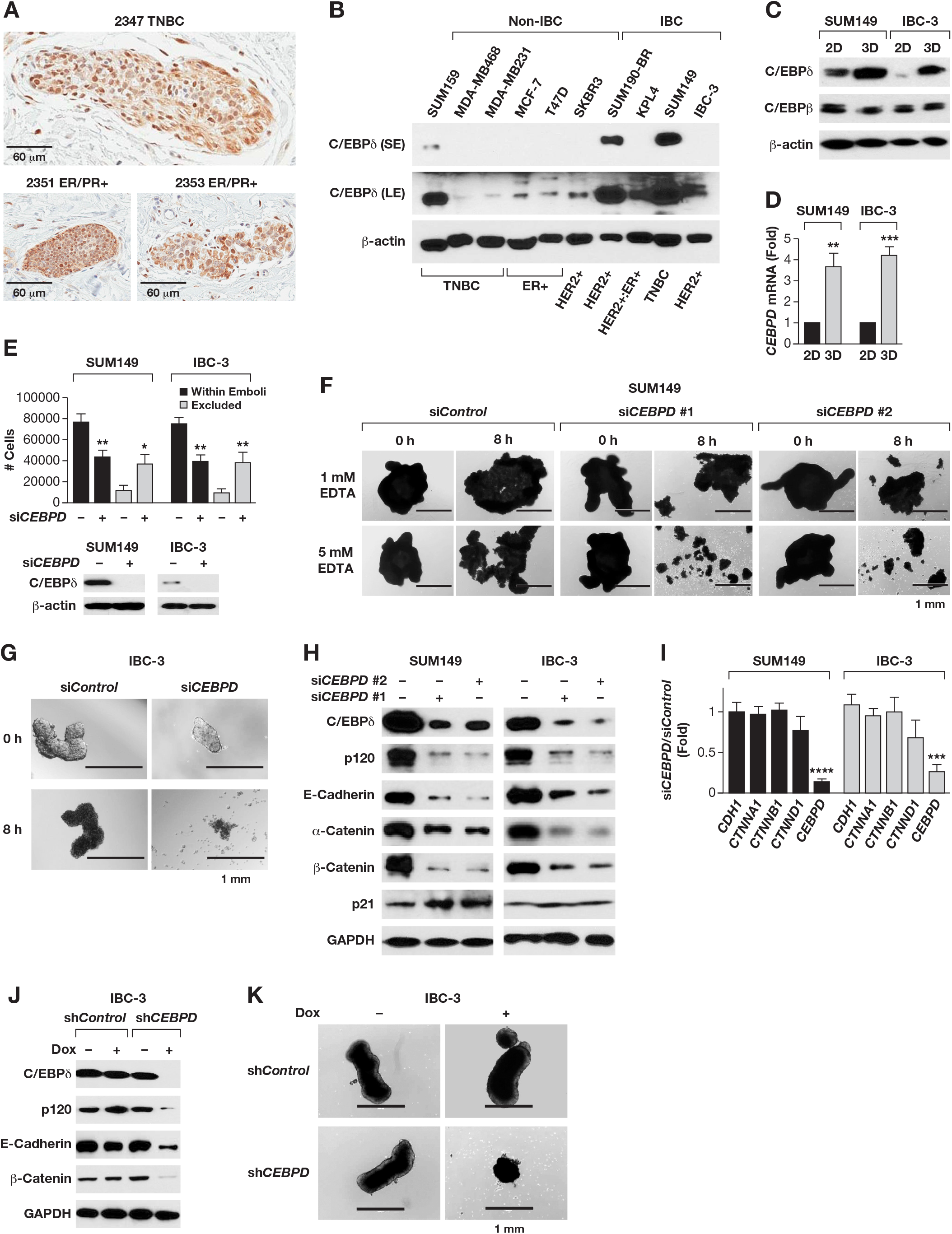
C/EBPδ is expressed in IBC emboli and IBC cell lines in vitro and promotes cell-cell adhesion and E-cadherin protein expression. **A**) C/EBPδ immunostaining in emboli from three independent IBC patient tissues. Scale bar=60 μm. **B**) Western analysis of C/EBPδ expression in whole cell extracts of the indicated cell lines (TNBC, triple negative breast cancer; ER, estrogen receptor α). β-actin was used as loading control. S/LE, short/long exposure. **C**) Western analysis of C/EBPδ and C/EBPβ in SUM149 and IBC-3 cell lines that were cultured on plastic (2D) or as emboli (3D) for 4 days. **D**) qPCR analysis of *CEBPD* mRNA in cells as in panel B (n=3, mean ±SEM; **P<0.01, ***P<0.001 compared to 2D). **E**) Quantification of SUM149 or IBC-3 cells transfected with si*Control* (-) or si*CEBPD* (+) oligos that aggregated into large clusters (“within emboli”) or remained as single cells / smaller clusters (“excluded”) after 3 days in 3D culture (n=3, mean ±SEM; \**P*<0.05, \*\**P*<0.01 compared to si*Control*). **F-G**) Images of similar-sized emboli from (F) SUM149 and (G) IBC-3 cells that had been transfected with control or two independent si*CEBPD* oligos before and after treatment with EDTA for 8 h (scale bar = 1 mm). **H**) Western analysis of the indicated proteins’ expression in established emboli of SUM149 and IBC-3 cells that had been transfected with siRNAs as indicated. **I**) qPCR analysis of *CDH1* (E-cadherin), *CTNNA1* (α-catenin), *CTNNB1* (β-catenin), *CTNND1* (p120) and *CEBPD* mRNA levels in emboli of SUM149 and IBC-3 cells transfected as in panel G (n=3, mean±SEM; ***P<0.001, ****P<0.0001 compared to si*Control*). **J**) Western analysis of IBC-3 cells with stable expression of the indicated shRNA and after culture in 3D for 3 days plus 3 days in the presence of doxycycline (Dox, 100 ng/ml). **K**) Images of representative emboli as in panel I and the same embolus before and after treatment with Dox (10 ng/ml) for 7 days (scale bar= 1 mm).

### C/EBPδ promotes expression of E-cadherin complex proteins through COX-2-mediated GSK3β inhibition

Because E-cadherin/catenin mRNA levels were not altered by C/EBPδ depletion, we tested whether C/EBPδ regulated protein stability. Treatment of emboli with the proteasome inhibitor MG132 increased E-cadherin, β−catenin, and p120 protein levels in *CEBPD*-depleted cells but had comparatively less effect on these proteins in control cells (**Fig. 2A**). E-cadherin protein stability depends in part on the formation of complexes at the cell membrane, which can be regulated by the abundance of β−catenin and p120 (21, 22), as was also demonstrated in SUM149 cells for p120 (22). The stability of β−catenin and p120 can be regulated by the serine/threonine kinase GSK3β which targets the proteins for degradation by the Skp1-Cullin-F-box^β-TrCP^ E3 ubiquitin ligase (21, 23). In C/EBPδ-depleted IBC cell lines, the inhibitory phosphorylation on Serine 9 (Ser9) of GSK3β (24) was significantly reduced, suggesting a higher level of GSK3β activity (**Fig. 2B** and **Fig. S2A**). p120, β−catenin and E-Cadherin protein levels were rescued in C/EBPδ-depleted IBC-3 cells when treated with two different GSK3β inhibitors, CHIR or LiCl, (**Fig. 2C**), and also when β-TrCP (*BTRC*) was silenced (**Fig. 2D**). Similar results were obtained with SUM149 cells (**Fig. S2B-C**). In control cells, GSK3β inhibition did not affect E-cadherin levels (**Fig. 2C, S2B**), suggesting that E-cadherin is not directly regulated by GSK3β. Notably, GSK3β inhibition significantly rescued the ability of *CEBPD*-silenced SUM149 and IBC-3 cells to associate into emboli (**Fig. 2E**). Taken together, these data show that C/EBPδ-mediated inhibition of GSK3β supports the accumulation of E-cadherin complex proteins and cell-cell adhesion.

**Figure 2.**
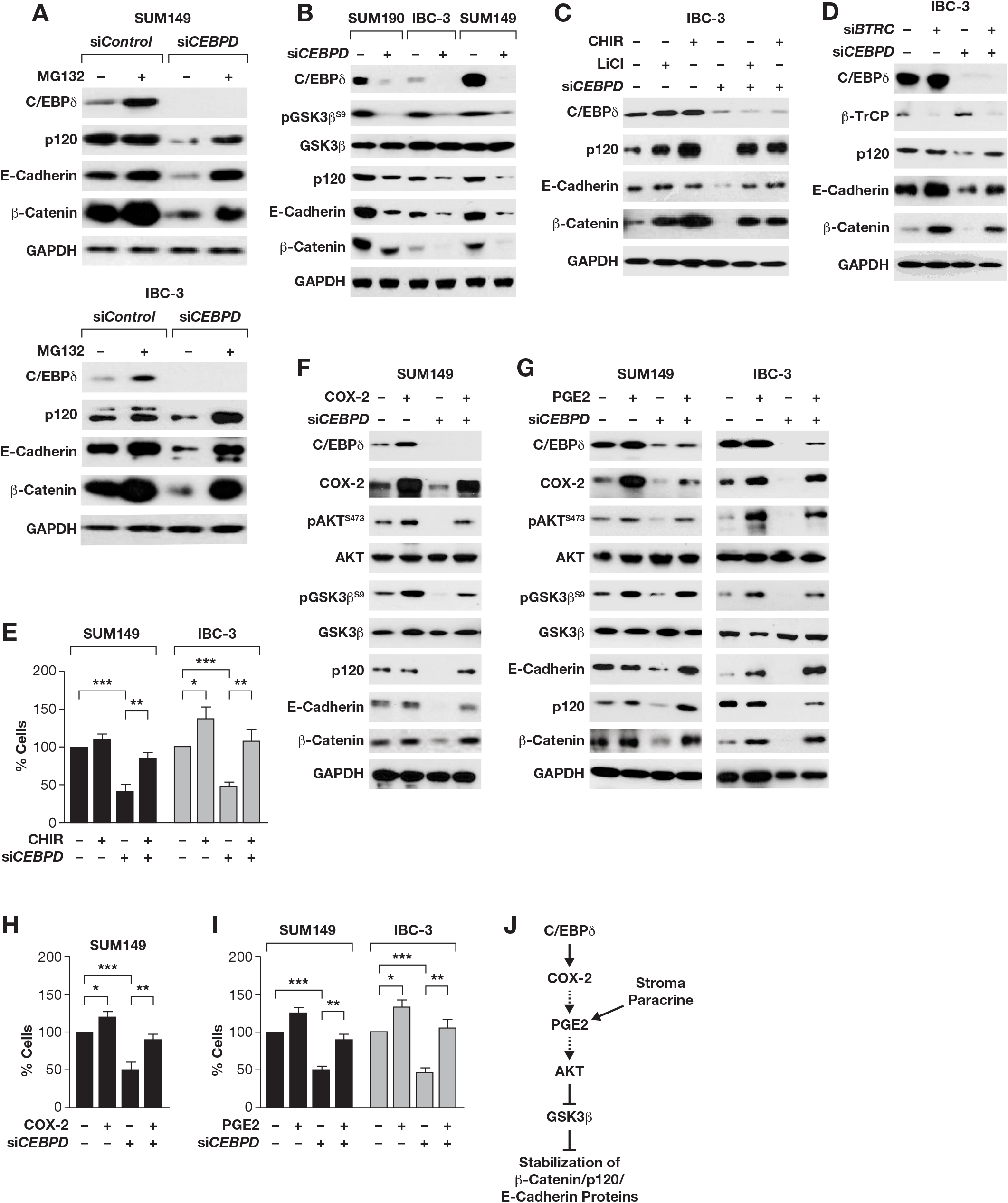
C/EBPδ promotes expression of E-cadherin complex proteins through COX-2-mediated GSK3β inhibition. **A**) Western analysis of emboli from SUM149 and IBC-3 cells transfected with siRNA as indicated and treated with 20 μM MG132 for 6 h. **B)** Western analysis of the indicated proteins in emboli from SUM190, IBC-3 and SUM149 cells that were transfected with control or si*CEBPD* oligos. **C)** Western analysis of the indicated proteins in emboli from IBC-3 cells transfected with control (-) or si*CEBPD* oligos and treated with LiCl (10 mM) or CHIR (5 μM) for 6 h. **D)** Western analysis of the indicated proteins in emboli from IBC-3 cells transfected with control (-) or si*CEBPD* along with si*BTRC* (β-TrCP) oligos. **E)** Analysis of the number of cells in emboli of SUM149 or IBC-3 cells that were transfected with si*Control* (-) or si*CEBPD* oligos and 24 h later seeded in 3D for 3 days ±1 μM CHIR (n=3, mean ± SEM, \**P*<0.05, \*\**P*<0.01, \*\*\**P*<0.001). **F**) Western analysis of the indicated proteins from SUM149 cells transfected with control (-) or si*CEBPD* (+) oligos and COX-2 expression plasmid followed by culture in 3D for 3 days. **G**) Western analysis of the indicated proteins in SUM149 and IBC-3 emboli by cells transfected as in panel A followed by culture in 3D for 3 days ±PGE2 (1 μM). **H**) Number of cells in SUM149 emboli as in panel F (% of control, n=3, mean ± SEM, \**P*<0.05, \*\**P*<0.01, \*\*\**P*<0.001). **I**) Number of cells in emboli of SUM149 and/or IBC-3 cells as in panel G (n=3, mean±SEM, \**P*<0.05, \*\**P*<0.01, \*\*\**P*<0.001). **J**) Model summarizing the signaling pathway described in this study and indicating that PGE2 may be generated by autocrine or paracrine/stromal mechanisms

Next, we investigated the mechanism by which C/EBPδ mediates GSK3β inhibition. GSK3β phosphorylation on Ser9 can be mediated by several kinases including AKT which is activated by many signaling pathways including that of prostaglandin E2 (PGE2), a downstream metabolite of the COX-2 enzyme. COX-2 is a target gene of C/EBPδ (25), correlates with AKT activation in BC (26, 27), and is highly expressed in IBC (28). Indeed, *CEBPD* silencing reduced COX-2 mRNA and protein expression in IBC cell lines (**Fig. S2D-E**), which was rescued by C/EBPδ overexpression (**Fig. S2F-G**). Along with downregulation of COX-2, *CEBPD* silencing also reduced AKT and GSK3β phosphorylation (**Fig. 2F**). Ectopic expression of COX-2 in C/EBPδ-silenced cells rescued phosphorylation of these proteins as well as expression of E-cadherin/catenin proteins (**Fig. 2F**). Similar results were obtained when *CEBPD*-depleted SUM-149 or IBC-3 cells were treated with PGE2 (**Fig. 2G**). Correspondingly, COX-2 overexpression (**Fig. 2H**) or PGE2 treatment (**Fig. 2I**) rescued the number of C/EBPδ-depleted SUM149 and IBC-3 cells associating into emboli. In summary, these data show that C/EBPδ-mediated COX-2 expression and activity lead to AKT activation and GSK3β inhibition in IBC cell emboli and that this pathway contributes significantly to the expression of epithelial cadherin complex proteins and cell-cell adhesion in 3D (**Fig. 2J)**.

### The COX-2/GSK3β/E-Cadherin pathway is conserved in a subset of breast cancers in vivo

To assess the potential in vivo relevance of our findings we examined COX-2 and E-Cadherin expression in clinical BC specimen. We focused our analyses on E-cadherin because it is the molecule that bridges cell-cell contacts. Co-occurrence of high COX-2 and E-cadherin expression (scores 3-4 for both) was observed in 48/172 (28%) of the breast tumors, was overrepresented in IBC (4/7 or 57%) compared to non-IBC (44/165 or 27%) (**Fig. 3A)**, and associated with worse BC-specific survival probability (**Fig. 3B**). Next, we evaluated E-cadherin and pGSK3β^S9^ in five metastatic PDX models, two ER+/PR+ (BCM-4888, -5097) models and three TNBC models (BCM-4013, -3204, -5471), with the latter expressing relatively more C/EBPδ (18). By immunostaining, all primary tumors as well as spontaneous PDX lung metastases, and SUM149 experimental metastasis, were positive for E-Cadherin and pGSK3β^S9^ (**Fig. 3C** and **Fig. S3A)**. Western blot analysis showed that all PDX models expressed COX-2, albeit at varying levels, and BCM-5471 the most (**Fig. 3D**), which was also seen at the level of mRNA (**Fig. S3B**). BCM-5471 also presented with local metastases that showed strong immunoreactivity for both E-cadherin and pGSK3β^S9^, such as a metastasis within a mammary duct or parenchymal tumor cell clusters resembling large emboli next to the primary tumor (**Fig. 3E-F**). Bronchial epithelial cells (**Fig. 3C**) and normal mammary epithelial cells (**Fig. 3E**) expressed high levels of E-cadherin, as expected, but were comparatively negative for pGSK3β^S9^. However, bronchial epithelial cells in close vicinity to metastatic lung lesions often exhibited stronger pGSK3^S9^ staining compared to more distal cells (**Fig. S3C**). While there can be other causes, this result is consistent with paracrine inhibition of GSK3β by factors such as PGE2. Collectively, these data indicate that co-expression of E-cadherin with pGSK3β^S9^ and COX-2 expression is observed in vivo in a subset of BCs including metastatic PDXs and could be indicative of aggressive tumor biology.

**Figure 3.**
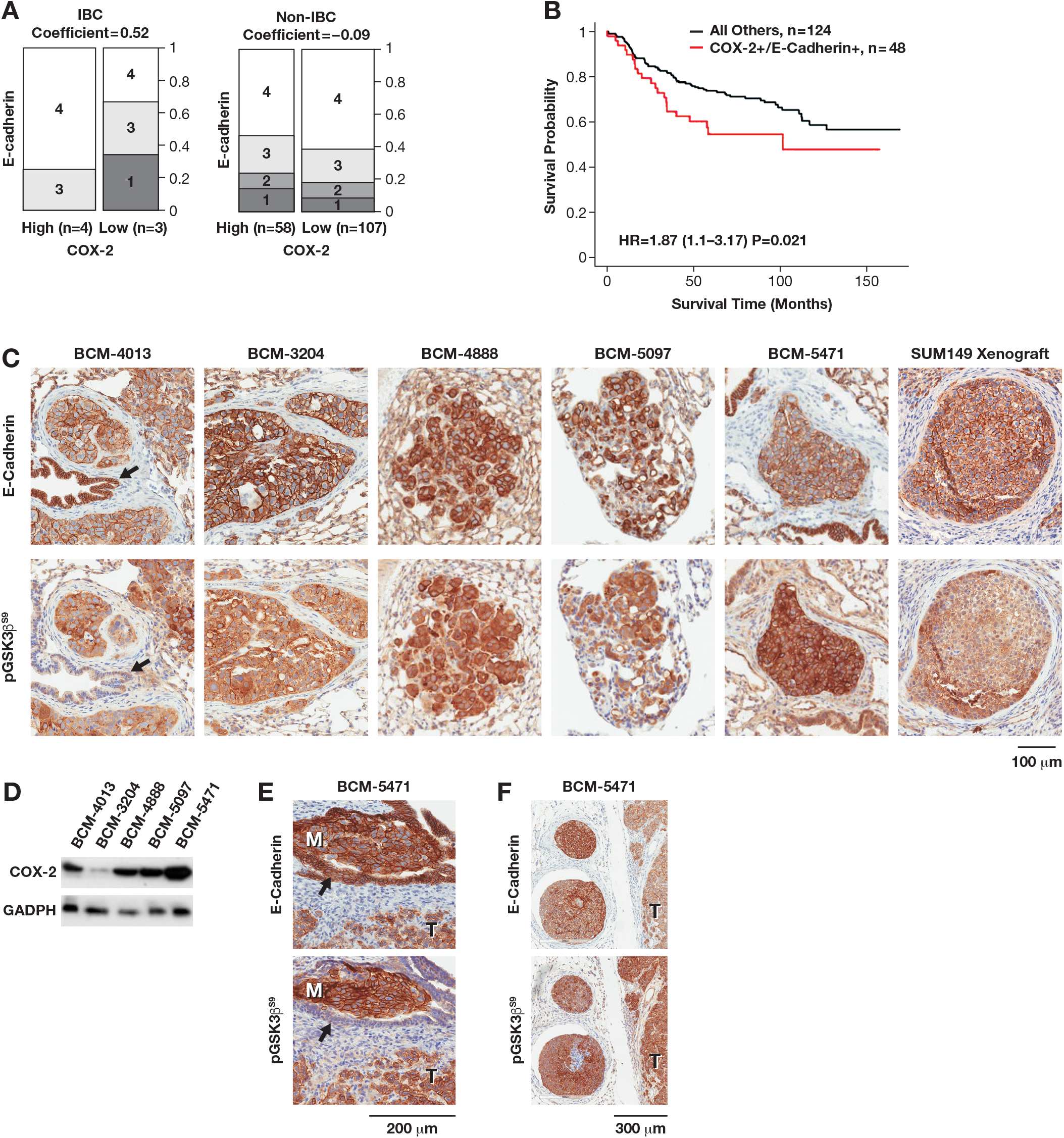
The COX-2/GSK3β/E-Cadherin pathway is conserved in a subset of breast cancers in vivo. **A**) Bar graph showing proportion of samples by different degrees of IHC staining of COX-2 and E-Cadherin in IBC (n=7) and non-IBC (n=165) tumor tissues. Numbers 1 to 4 within boxes (along with dark to lighter shades of grey) denotes low to high expression levels of E-cadherin. Columns represent high (score 3-4) versus low (score 1-2) COX-2 expressing samples. Width of columns and scale denotes relative proportion of samples with different combinations of scores. “Coefficient” refers to Pearson correlation coefficient for COX-2 and E-cadherin expression. **B**) Kaplan-Meier plot with the hazard ratio (HR) and 95% confidence interval from a Cox regression analysis comparing COX-2 high/E-cadherin high patients with all other patients (reference group). Patients with high COX-2 and high E-cadherin expression (COX-2+/E-cadherin+) in their tumors have a significantly decreased breast cancer-specific survival when compared with all other patients (P=0.021). **C**) Immunostaining of E-Cadherin and pGSK3β^S9^ on serial sections of lung metastases from PDX primary tumors, representing TNBC (BCM-3204, -4013, 5471), ER+/HER2+ (BCM-4888), and ER+ (BCM-5097) subtypes, and an experimental metastasis by SUM149 cells. Black arrow indicates bronchial epithelium (BCM-4013). **D)** Western analysis of COX-2 in tumor tissue extracts from the indicated PDX models. **E)** Immunostaining as in panel C of BCM-5471 showing a micrometastasis within a mammary duct (M, metastasis; T, tumor; arrow, mouse mammary epithelium). **F)** BCM-5471 as in panel C showing emboli-like structures next to primary tumor, T.

### The COX-2 inhibitor celecoxib downregulates E-cadherin protein in vivo and reduces SUM149 tumor growth and circulating tumor cells

To determine the effect of pharmacological COX-2 inhibition on E-cadherin/catenin expression, we treated established in vitro emboli with celecoxib, which inhibited AKT and GSK3β phosphorylation within 24-48 h, along with reducing expression of β−catenin, p120 and E-cadherin (**Fig. 4A)**. Celecoxib also downregulated COX-2 and C/EBPδ expression, consistent with autoregulation respectively positive feedback regulation, while p21^CIP1/WAF1^ expression was induced (**Fig. 4A)**. These events were followed by induction of cell death (**Fig. 4B** and **Fig. S4A**). When added at the time of seeding in 3D, celecoxib prevented cell aggregation and some cells underwent cell death by day 3 (**Fig. 4C**). Celecoxib also downregulated ectopic E-cadherin protein (**Fig. 4D**), which confirms that COX-2 supports E-Cadherin expression at the protein level and explains why ectopic E-cadherin could not rescue cell survival in 3D **(Fig. 4D**).

**Figure 4.**
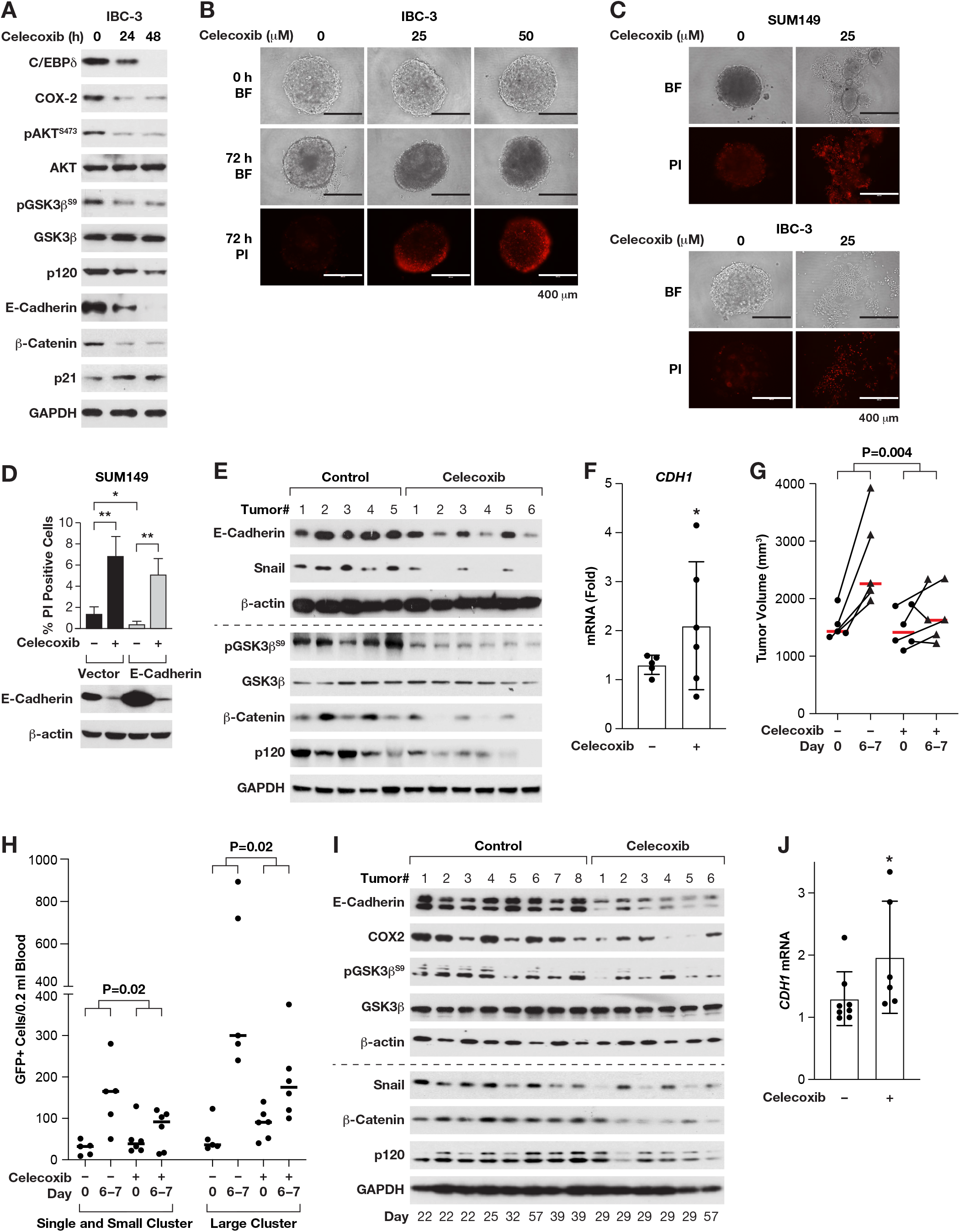
The COX-2 inhibitor celecoxib downregulates E-cadherin protein in vivo and reduces SUM149 tumor growth and circulating tumor cells. **A**) Western analysis of IBC-3 emboli established after 3 days of culture in 3D followed by treatment for the indicated times with 50 μM celecoxib (0h = 48 h DMSO). **B**) Images of representative IBC-3 emboli after 3 days of culture (0 h) and the same emboli following another 72 h with celecoxib and stained with propidium iodide (PI) to label dying cells as indicated (BF, bright field; Scale bar = 400 μm). **C**) Representative images of SUM149 and IBC-3 cells cultured in 3D ± celecoxib for 72 h and stained with PI (BF, bright field; Scale bar = 400 μm. **D**) Assessment of cell death by PI staining (top panel) and Western analysis (bottom panel) from SUM149 cells that were transfected with empty vector or E-Cadherin expressing plasmid followed by culture in 3D for 1 day and treated with CXB for additional 3 days (n=3, mean±SEM. *P>0.05, **P>0.001). **E-G**) Western (E), *CDH1* mRNA (F) and tumor volume (G) analysis of SUM149-GFP-Luc orthotopic tumors from mice fed control chow or celecoxib-chow for 6-7 days starting at tumor volumes >1000 mm^3^, n=5-6 (panel F, *P<0.05; panel G, P=0.004 by unpaired Wilcoxon test (larger increase in tumor volume over time for untreated versus treated mice). **H**) CTC analysis of peripheral blood drawn from mice as in panels E-G (n=5-6, P=0.023 by Linear Model of Day-Treatment Interaction between groups of treated versus untreated mice). **I-J**) Western (I), and *CDH1* mRNA (J) analysis of BCM-5471 PDX tumors from mice that were fed control chow or celecoxib-chow for the indicated number of days (determined by study endpoints) starting when tumor volumes were 300-600 mm^3^ (n=6-8, panel J: P=0.029 by unpaired two-sided Wilcoxon rank-sum test).

Next, we evaluated the effect of celecoxib on SUM149 orthotopic primary tumor xenografts. Treatment of mice with large tumors for 6-7 days downregulated expression of the EMT factor Snail (**Fig. 4E**), as described previously in gastric cancer models (29). Nonetheless, tumors of celecoxib-treated mice expressed significantly less E-cadherin complex proteins compared to untreated (**Fig. 4E**), although *CDH1* mRNA levels were modestly increased (**Fig. 4F**). Despite the downregulation of E-cadherin protein, the tumor growth rate was attenuated by celecoxib (**Fig. 4G**). Quantification of circulating tumor cells (CTCs) through expression of a GFP reporter before and after celecoxib treatment demonstrated that their numbers increased over time in untreated mice but not in treated mice (**Fig. 4H**). As an alternate non-IBC model system, we also treated BCM-5471 PDX tumors with celecoxib and again observed that E-cadherin expression was reduced at the level of protein but not mRNA **(Fig. 4I-J)**, along with reduced levels of COX-2, pGSK3β^S9^, β−catenin, and p120 (**Fig. 4I** and **Fig. S4B**). Taken together, these data from 3D culture and two in vivo model systems show that a therapeutic potential of celecoxib is accompanied by downregulation of E-cadherin protein expression in vivo.

### Celecoxib cooperates with paclitaxel in attenuation of experimental and spontaneous lung metastases

Lung metastases initiate as intravascular emboli that require E-cadherin as has been shown through antibody-based inhibition (6). We corroborated this notion by a genetic approach in which the E-cadherin gene was deleted by inducible Cre-recombination in mouse mammary tumor cells (12) after the onset of lung colonization, which significantly reduced the tumor burden in lungs **(Fig. S5A)**. Despite the presence of CTCs (**Fig. 4H**), in our hands, only about 10% of mice with SUM149 xenografts developed spontaneous lung metastases. Thus, we proceeded to evaluate experimental lung metastases generated after tail vein injection of luciferase expressing SUM149 cells. When bioluminescence imaging (BLI) confirmed lung colonies, mice were randomized to treatments. We used not only celecoxib but also paclitaxel after having determined the combinatorial benefit of these two drugs in 3D culture (**Fig. S5B-C**). At dosing as previously reported for combination treatments (30, 31), both paclitaxel and celecoxib monotherapy reduced BLI signal in the lungs compared to untreated mice while the combination therapy completely eliminated bioluminescence (**Fig. 5A)**, which was confirmed by histological evaluation of lungs (**Fig. S5D**). When the doses were halved, monotherapies were no longer effective, but combination therapy significantly attenuated the BLI signal (**Fig. 5B)**. These results show that celecoxib can diminish established SUM149 experimental lung metastases and synergizes with paclitaxel treatment. The data also indicate that the 3D emboli culture paradigm modeled SUM149 cell responses in vivo. Next, we proceeded to evaluate the drug response of BCM-5471. Because this PDX model does not express a luciferase reporter, we began treatment when tumors were well established and likely to have seeded lung metastases. Celecoxib alone and combination treatment slowed primary tumor growth to varying degrees (**Fig. S5E**) and the combination treatment resulted in reduced tumor volumes after 22 days of treatment (**Fig. 5C**). Histological quantification of spontaneous micrometastases showed that the monotherapies had no significant effect but that the lungs of mice under combination treatment harbored significantly fewer tumor cells than untreated mice (**Fig. 5D-E**). Taken together, these data demonstrate a therapeutic benefit of celecoxib alone (SUM149) or in combination with paclitaxel (SUM149, BCM-5471) in reducing both experimental and spontaneous lung metastases by cells expressing both COX-2 and E-cadherin.

**Figure 5.**
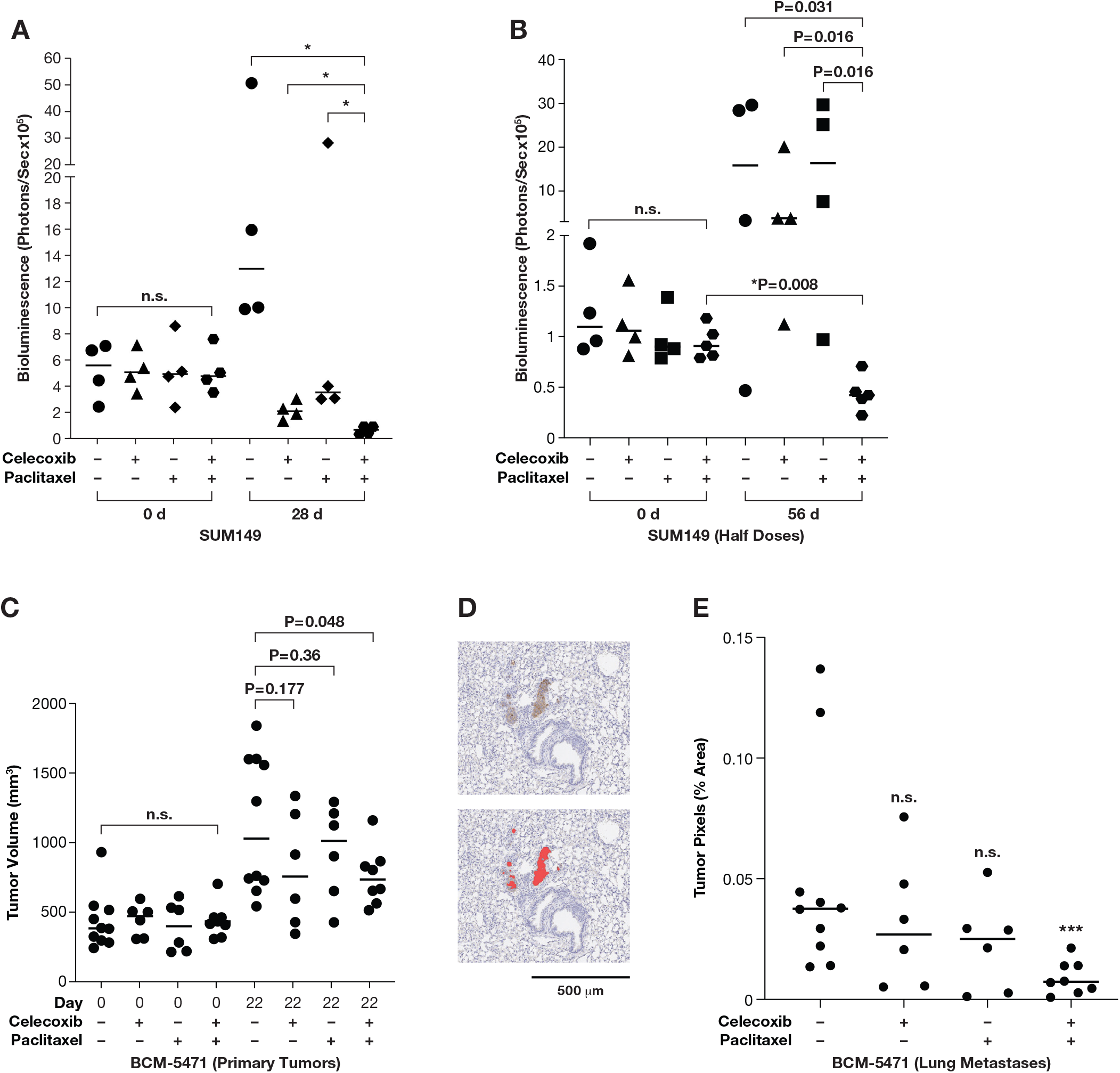
Celecoxib combination with paclitaxel attenuates experimental and spontaneous lung metastases. **A)** Quantification of bioluminescence in the lungs of mice (n=4) with experimental metastases of SUM149-GFP-Luc cells before (day 0) or after 28 days of treatment with celecoxib 1000 mg/kg chow) and/or paclitaxel (10 mg/kg i.v.). *P=0.028 by unpaired two-sided Wilcoxon rank-sum test, n.s. indicates no significant difference between any of the groups on day 0. **B)** Quantification of bioluminescence in mice (n=4-5) as in panel A after 56 days of treatment with celecoxib (500 mg/kg chow) and/or paclitaxel (5 mg/kg i.v.). P values as indicated by unpaired or paired (*P) two-sided Wilcoxon rank-sum test; n.s., as in panel A. **C**) Tumor volume measurements of BCM-5471 PDX in mice on day 0 and 22 of treatment as in panel A (n=6-10, P values as indicated by two-sided t-test. **D)** Light microscope image of a mouse lung section from the experiment in panel C showing representative micrometastases immunostained with human-specific “mitomarker” (top panel) and their pixilation by the Halo image analysis software (bottom panel). **E**) Quantification of tumor cell pixels in representative sections of lungs from mice as in panel C. % of total lung area, n=6-10, ***P=0.0005 by Unpaired Wilcoxon test.

## DISCUSSION

In this study, we have mechanistically connected two seemingly disparate aspects in cancer biology: the role of inflammation in BC metastasis as exemplified by COX-2 signaling and expression of E-cadherin mediating cell-cell adhesions. We show that COX-2 and E-cadherin are co-expressed in clinical specimen with poor survival probability and metastatic TNBC PDX models. Furthermore, the COX-2 inhibitor celecoxib downregulated E-cadherin while also attenuating primary tumor growth and reducing CTCs and, especially when combined with paclitaxel, experimental and spontaneous lung metastases. In addition to promoting cell-cell adhesion that can foster cell survival under adverse conditions, elevated levels of E-cadherin expression could also lead to increase in soluble extracellular or cytoplasmic fragments that have oncogenic properties (32, 33). Through studies of IBC cells and their ability to form “emboli” in culture, we resolved a molecular mechanism by which COX-2 signaling supports E-cadherin protein expression via GSK3β inhibition, possibly through direct stabilization of p120 and β-catenin. A variety of 3D culture paradigms have been established and shown to mimic more closely the physiological contexts compared to cell culture on plastic (34). The most unique feature of “emboli culture” compared to other 3D culture paradigms is the mechanophysical environment (14). Our ongoing studies are addressing to what extent specific 3D culture methods affect signaling pathways. However, in the current report, we demonstrate that the “emboli culture” method replicates the effect of COX-2/PGE2 signaling seen in vivo, i.e. inhibition of GSK3β and supporting expression of E-cadherin complex proteins. Future studies will need to address if and how this pathway may be enriched in IBC compared to non-IBC.

We identified C/EBPδ as a tumor cell intrinsic factor that can initiate the COX-2/E-cadherin pathway. C/EBPδ is most highly expressed during the first, inflammatory phase of postpartum mammary gland involution (35) and again in the fully involuted stage (36). These conditions, which also involve COX-2 signaling, promote the risk of aggressive post-partum BC including IBC (37-39). However, COX-2 can also be induced by C/EBPδ-independent pathways, and PGE2 can also be provided through tumor cell extrinsic microenvironmental sources (37). Indeed, stromal expression of COX-2 expression was seen in canine IBC compared to non-IBC (40, 41). Thus, in addition to tumor cell intrinsic C/EBPδ and COX-2, our results provide a molecular mechanism by which E-cadherin expression can be maintained in metastatic BC cells through PGE2 and/or inhibition of GSK3β. Many oncogenic kinases including AKT can inhibit GSK3β (24) but are not necessarily associated with E-cadherin expression. Similarly, many studies reported on a role of COX-2 in promoting EMT and that COX-2 inhibition prevents EMT (42-44). Our results demonstrate that E-cadherin (*CDH1*) mRNA levels do not always accurately reflect E-cadherin protein expression and that epithelial-mesenchymal plasticity can be attained through stabilization of E-cadherin complex proteins. In addition, the difference in results between these studies and ours may be in part due to culture conditions and/or cell type and context. In BC specifically, the intrinsic subtypes represent “unique diseases” (45). We obtained comparable results with two different IBC cell lines in vitro (TNBC SUM149, HER2+ IBC-3), and a TNBC cell line (SUM149) and PDX model (BCM-5471) in vivo. Future studies will have to determine to what extent the context of BC subtype within ductal carcinomas may modulate the relationship of COX-2 signaling, E-cadherin protein expression, and metastasis.

Both, antibody-based inhibition (6, 46) and genetic deletion reveal a critical role for E-cadherin in promoting cancer cell survival and metastasis in invasive ductal BC ((12, 47) and this study). Furthermore, knockdown of E-cadherin impaired the growth of primary tumors and experimental metastases in TNBC xenograft models (11) and in a luminal GEMM model (12). These reports corroborate our observation that downregulation of E-cadherin protein by celecoxib was accompanied by attenuated tumor growth and reduction in CTCs and lung metastases with increased sensitivity to paclitaxel treatment. Combination therapies with celecoxib and including taxanes have been tested in preclinical models and clinical trials and showed some indication of efficacy (48). For successful COX-2-targeted therapies of breast cancer, however, these trials revealed an unmet need to better identify which patients will respond (49, 50). While downregulation of E-cadherin may not always be necessary for response to celecoxib, our study showed that COX-2 can maintain cell-cell adhesions in ER-negative aggressive breast cancer cells through GSK3β inhibition and suggests that combined evaluation of E-cadherin protein, COX-2, and pGSK3β^S9^ could potentially contribute to the identification of patients with metastatic BC who may benefit from combination therapies that target the PGE2 signaling pathway.

## MATERIALS AND METHODS

### Antibodies

Antibodies were obtained from the following sources, unless indicated otherwise: Cell Signaling Technology (pSTAT3^Y705^, #9145; STAT3, #4904; Cleaved Notch-1, NICD, #2421S; E-cadherin, #3195 (24E10), #5296 (32A8), α-catenin, #3236S; β-catenin, #9562S; COX2, #12282S; pGSK3β^S9^, #9323T; GSK3β, #9315S; pAKT^S473^, #9271S; AKT, #4691; p21, #2947S; Snail, #3879S); eBiosciences (CD44-PE, #12-0441-82, clone IM7; CD24-FITC, #11-0247-41, clone eBioSN3; CD24-APC, #17-0247-42, clone eBioSN3), Abcam (β-actin, ab6276 (Ac-15); Santa Cruz Biotechnology (CXCR4, #sc-9046; GAPDH, #sc-47724); Rockland (α-tubulin, #600-401-880), BD Biosciences (p120, #610133).

### Cells, culture and reagents

MCF-7, T47D, MDA-MB-468, MDA-MB-231, and SKBR3 cells were obtained from ATCC; SUM149 and SUM190 cells originated from Asterand Bioscience. IBC-3 and KPL-4 cells were kindly provided by Dr. Wendy A. Woodward (MDACC) and Dr. Junichi Kurebayashi (Kawasaki Medical School), respectively, brain tropic SUM190-BR cells by Dr. Patricia Steeg (NCI), and SUM149 derivatives by Dr. Jangsoon Lee (MDACC) and Dr. Stanley Lipkowitz (NCI). Cell lines were authenticated in 2014 and/or 2017 and/or 2019 by GenePrint®10 (Promega), and *Mycoplasma* tested about annually by qPCR. Cells were cultured in a 5% CO_2_ incubator at 37°C in media with 100 units/ml penicillin and 100 μg/ml streptomycin as follows: MCF-7, MDA-MB-231, and MDA-MB-468 in Dulbecco’s modified Eagle’s medium (DMEM), MCF-7 also with 1 mM sodium pyruvate; T47D in ATCC-formulated RPMI-1640 Medium (#30-2001) with 0.2 units/ml bovine insulin (Sigma-Aldrich, #I0516); SUM159 in RPMI with 2 mM glutamine, 10 mM HEPES, 1 mM sodium pyruvate, 1X nonessential amino acids (GIBCO, #11140-050) and 55 μM β-mercaptoethanol (GIBCO, #21985-023); SKBR3 cells in McCoy’s 5A Medium Modified (GIBCO, #16600-082); SUM149, IBC-3 and SUM190 in Ham’s F-12 media (GIBCO, #31765092_)_ with 1 μg/ml hydrocortisone and 5 μg/ml Insulin; KPL-4 cells in DMEM/F12/GlutaMax™. Fetal bovine serum (FBS) was added at 10% except for SUM159 (5%). Cell culture grade chemicals were from Sigma Aldrich unless indicated otherwise.

Celecoxib (#NDC 59762-1517-1) and paclitaxel (#NDC-0703-3213-01) were purchased from the NIH Pharmacy (Bethesda, MD); carboplatin (#S1215), Prostaglandin E2 (#S3003), and CHIR-99021 (#S2924) were from Selleck Chemicals, USA; Doxorubicin (#D1515) and propidium iodide (#P-4170) were from SIGMA. DMSO (SIGMA, #D-2650) was used as vehicle control in all experiments.

### 3D culture assay

*In vitro* emboli formation was carried out as described (14). Briefly, cells were trypsinized 24 h after nucleofection (if applicable), 100,000 cells were seeded in 6-well ultra-low attachment plates (Corning, #3471) in medium containing 2.25% PEG8000 and gently rocked at approximately 40 rpm for 3-4 days or as indicated. To isolate emboli, cultures were centrifuged at 500 rpm for 1 min with PBS, treated with TrypLExpress (GIBCO, #12604-013) for 5-10 min and neutralized with cell culture medium. Cells were counted with a Countess (ThermoFisher) using trypan blue dye exclusion. Unless indicated otherwise, all analyses of emboli were conducted after 4 days in 3D culture. For assessment of cell death within established emboli, these were generated first by seeding 10,000 cells per well in 96-well plates (Nexcelom Biosciences, Cat#ULA-96U-010), cultured and treated as indicated, followed by addition of propidium iodide (PI, 0.5 μg/ml) for 30 min and imaging with an EVOS® FL microscope. For quantification, emboli were harvested as above, cells transferred at 10,000 cells/well in 96-well plates and 6 h later treated with PI for 30 min and analyzed by Direct Cell Counting (Celigo, Nexcelom).

### Invasion assay

Cellular invasion through Matrigel was carried out using Corning BioCoat-growth factor reduced 24 well plates according to the manufacturer’s protocol (Corning, U.S., # 354483). Briefly, SUM149, IBC-3 and KPL-4 cells were nucleofected with control or *CEBPD* siRNA oligos. Seventy-two hours later 5 × 10^4^ cells in serum-free medium were placed in the chamber and immersed in 24 well plates with serum-containing medium and incubated at 37°C for 8 h. After fixing with 4% formaldehyde for 2-5 min followed by methanol for 10-15 min, the cells were stained with crystal violet for 15 min. Migrated cells on the entire surface of the membrane were viewed under the microscope and counted manually and blinded to the experiment.

### Flow cytometric analysis

About 2×10^5^ cells per sample were blocked using Purified NA/LE Rat Anti-mouse CD16/CD32 Clone 2.4G2 antibody (BD Biosciences, #553140) followed by incubation with 1 μl of specific antibodies for 30 min on ice in the dark. Isotype specific antibodies (see Antibodies) and/or OneComp eBeads (eBiosciences, #01-1111-42) were used as negative controls. Cells were washed twice with ice-cold PBS containing 0.02% sodium azide, resuspended in DPBS/0.1% BSA and analyzed with a BD FACSCanto II Analyzer and FlowJo software (FlowJo, LLC., Ashland, OR). At least 30,000 viable events per sample were collected for analysis.

### Generation of cells with stable or Dox-inducible shRNA expression

For stable shRNA expression, SUM149 cells were infected with pDEST lentiviral vector expressing sh*CEBPD* or *GFP*-targeting (GCAAGCTGACCCTGAAGTTCAT) sh*Control* RNA, packaged with MISSION Packaging Mix (SIGMA cat# SHP001), and selected by G418. SUM149 and IBC-3 cells with Dox-inducible shRNA expression were first infected with CMV-Luciferase-2A-GFP (Neo) (GenTarget Inc, Cat# LVP403) virus and selected as per instructions. Subsequently, cells were infected with SMARTchoice lentivirus from Dharmacon (Non-targeting control, Cat# VSC11656; CEBPD shRNA, Cat# V3SH11252-226035621) and selection per instructions. For shRNA induction, cells were treated with doxycycline as indicated. The sequences of the siRNA and/or shRNA used in this study can be found in Supplementary **Fig. S6**.

### Transient expression and silencing of gene expression

pcDNA3.1-hPTGS2-2flag(51) was a gift from Dr. Jun Yu (Addgene plasmid # 102498; http://n2t.net/addgene:102498 ; RRID:Addgene_102498). siRNA-mediated silencing was by nucleofection with AMAXA^®^ technology essentially as described (52). All experiments included non-specific siRNA (-) as control. Scrambled siRNA was used for most experiments (sense 5′-CGUACGCGGAAUACUUCGAUUdTdT-3′), *GFP* oligonucleotides were used alternatively (sense 5′-CAAGCTGACCCTGAAGTTC-3′). Unless indicated otherwise, *CEBPD* siRNA#1 (sense 5’-UCGCCGACCUCUUCAACAGTT-3’) and *CEBPD* siRNA#2 (sense: 5’ -CCACUAAACUGCGAGAGA-3’) were used at 1:1 ratio. *BTRC* siRNA: Sense 5’-GUGGAAUUUGUGGAACAU-3’. For each experiment, the efficiency of silencing was assessed by Western and/or qPCR analysis.

### Western analysis

Whole cell extracts were prepared by lysing the cells or emboli with RIPA buffer (25 mM Tris, pH 8.0, 50 mM NaCl, 1 mM EDTA, 0.5% NP40, 0.5% sodium deoxycholate, 0.1% SDS; 10 μl/ml protease inhibitor cocktail, Sigma #P8340; 10 μl/ml phosphatase inhibitor cocktail #2, Sigma #P5726; 10 μl/ml phosphatase inhibitor cocktail #3, Sigma #P0044). Tumor extracts were prepared as described (18). Protein concentrations were measured by BCA assay (Thermo Fisher, cat#23225). About 10-20 μg protein was loaded onto NOVEX WedgeWell 4-20% Tris-glycine gels and Western analyses were carried out as described (52).

### RNA isolation and quantitative PCR

Total RNA from cell lines and tumor tissues was purified by GeneJET RNA purification kit (Thermo Scientific, #K0732), and cDNA was synthesized using Superscript Reverse Transcriptase III (RT) according to manufacturer’s instructions (Invitrogen, #18080044). PCR was carried out with Fast SYBR Green master mix (#4385612, Applied Biosystems, Foster City, USA) using the 7500 Fast Real**-**Time PCR instrument (Applied Biosystems) and the relative expression levels were measured using the relative quantitation ΔΔ*C*t method and normalized to *RPLP0*. Data are from three independent biological replicates, each assayed as triplicates. For the primer details, see Supplementary **Table S1**.

### PDX and mice

Tissue sections of primary tumors (transplant generation 6-13) and lungs were obtained from the NCI-CCR Breast Cancer PDX Biobank. All PDX models were previously established and characterized at Baylor College of Medicine, https://pdxportal.research.bcm.edu/pdxportal (53). BCM-5471 was propagated in NOD/SCID/ILIIrg−/− (NSG) mice essentially as described (53). Experiments were performed with transplant generations 9-10. Tumor volumes were calculated as V = (W(2) × L)/2. Celecoxib (#NDC 59762-1517-1, NIH Pharmacy, Bethesda, MD) was provided in powder feed (AIN-93G, Envigo) at 1000 mg/kg chow as described (30). Paclitaxel (NDC 47781-593-07, NIH Pharmacy, Bethesda, MD) was administered i.v. at 10 mg/kg, once a week. Ground chow without celecoxib and injection of vehicle (50:50 ethanol:Kolliphor® to 5 parts saline) were used as controls. Treatments were started when tumors were established (300-800 mm^3^) and for the indicated durations.

### SUM149 xenografts and CTC analysis

SUM149-GFP-Luc cells (3×10^6^) were injected into the inguinal fat pads of 9-17 week old female NSG mice. When tumors reached about 1100-1900 mm^3^ volume, treatment started with celecoxib (1000 mg/kg of chow) or normal powder feed for 6-7 days. About 200 μl blood was collected from the tail vein before and after the treatment. Erythrocytes were lysed using ACK lysing buffer (Lonza, #BP10-548E) and remaining cells were washed with PBS and suspended in 200 μl PBS, seeded into 2 wells (Corning#655090) and analyzed for GFP+ cells using Celigo Imaging Cytometer. Gating was designed for 3 different populations: < 201 μm^2^ area to denote debris or small single CTCs; 201-400 μm^2^ to label single and small cluster GFP+CTCs; >400 μm^2^ area to denote large cluster CTCs.

### Experimental metastasis assays

SUM149-GFP-Luc cells (2-3×10^6^ in PBS) were injected i.v. into 6-8 week-old female nu/nu mice. Mice were monitored biweekly for bioluminescence and randomized by bioluminescence intensity for treatment with celecoxib (1000 mg/kg chow) and paclitaxel (10 mg/kg) 5 times/10 days plus 2 weekly doses (for data in Figure 5A) or with celecoxib (500 mg/kg chow) and paclitaxel (5 mg/kg) once per week for 8 weeks (for data in Figure 5B). In vivo bioluminescence imaging (IVIS Spectrum imager, PerkinElmer Inc.), was performed essentially as described (30). Bioluminescence signals were quantified by Living Image (version 4.3.1, PerkinElmer Inc.), implementing standard region of interests (ROI) drawn over the metastatic region. MMTV-PyMT mouse mammary tumor cells with E-cad^fl/fl^ or E-cad^fl/fl^; CreER as well as mTomato and mGFP as described (12) were injected as small clusters (about 2×10^5^ cells) into 6-8 week old NSG mice (12). One week later, all mice were injected with tamoxifen (100 μl of a 2 mg/ml stock) to delete E-cadherin and induced mGFP expression in cells with E-cad^fl/fl^; CreER cells. Three weeks later, lungs were harvested and the number of metastases as red and/or green-fluorescent foci were counted blinded under a dissection microscope. These experiments were performed in accordance with protocols approved by the Johns Hopkins Medical Institute Animal Care and Use Committee (ACUC).

### Histological analysis of tumor and lung tissue

Immunohistochemistry was performed with primary antibodies for E-cadherin at 1:400 (Cell Signaling Technology #3195), pGSK3β^S9^ at 1:100 (Abcam #75814) and C/EBPδ at 1:100 (Santa Cruz, sc-135733) with relevant positive and negative controls. For quantification of SUM149 lung metastases, four 5 μm sections, 100 μm apart, were stained with hematoxylin and eosin and evaluated by a veterinarian pathologist blinded to the experiment. For the quantification of PDX lung metastases, sections were stained with human-specific anti-mitochondria antibody (Abcam, ab79479), and scanned slides were analyzed with Halo-imaging software to quantify tumor cell area per total lung area of the most representative section.

### Patient survival analysis

IHC of COX-2 and pAKT protein expression in 248 human breast tumors (17 IBC, 58 TNBC, and 42 HER2+ and 145 ER+ and 102 ER-negative tumors) was previously reported (26). This research has previously been approved by the NIH Office of Human Subjects Research Protections (OHSRP #2248) and followed the ethical guidelines set by the Declaration of Helsinki. IBC samples, 6 TNBC and 1 HER2+, were classified as described (54). E-cadherin IHC (Dako M3612 antibody at 1:100) was available for 172 of these tumors (7 IBC, 42 TNBC, 31 HER2+, 98 ER+, and 73 ER-negative). Protein expression in the tumor epithelium was scored as negative, low, moderate, or high, and then categorized into low (negative to low) and high (moderate to high) for correlation and survival analysis, as previously described (26). We performed a Pearson correlation test to evaluate relationships between protein marker expression and tumor characteristics. We used Cox proportional hazards regression to estimate hazard ratios (HRs) with 95% confidence intervals (CIs) to assess the association between marker expression and BC survival. Survival curves were generated using Kaplan-Meier plots.

### C/EBPδ immunostaining in IBC patient tissues

IBC tissues were drawn from the IBC registry at MDACC as described (55). One section per specimen was stained from mastectomies of 39 patients clinically characterized as IBC and who had not achieved complete pathological response after primary systemic treatment. The specimen represented the following subtypes: 25 ER+/HER2-, 3 ER+/HER2+, 2 ER-/HER+, 9 TNBC, 1 undefined. Of the analyzed sections, 14 specimens presented with emboli in the tumor parenchyma. The data analysis for this research was approved by the Institutional Review Board of the MD Anderson Cancer Center. Immunohistochemistry of C/EBPδ was performed as described with monoclonal antibody 92.69 (19).

#### Statistics

Unless stated otherwise, quantitative data were analyzed by the two-tailed unequal variance t-test and are shown as the mean±S.E.M. The number of samples (n) refers to biological replicates.

#### Study approval

Research on patient material has previously been approved by the NIH Office of Human Subjects Research Protections (OHSRP #2248) and followed the ethical guidelines set by the Declaration of Helsinki. For studies with animals, NCI-Frederick is accredited by AALACi and follows the Public Health Service Policy for the Care and Use of Laboratory Animals. Animal care was provided in accordance with the procedures outlined in the “Guide for Care and use of Laboratory Animals” (National Research Council, 1996; National Academy Press; Washington, D.C.). All experiments were conducted under protocols approved by the Animal Care and Use Committee, NCI-Frederick, and in accordance with the NIH Guide for the Care and Use of Experimental Animals.

## Author Contributions

Designed research (KB, ES), conducted experiments and acquired data (KB, S.Se, S.Sh, LM, VP, KC), analyzed data (KB, S.Se., SK, WT, VP, SA, AJE, DKP, ES), provided reagents (SK, AJE, NTU, SA), wrote manuscript (KB, DKP, ES), edited manuscript (KB, SK, AJE, NTU, SA, DKP, ES).

## ACKNOWLEDGEMENTS

We are grateful for superb support by the NCI/CCR Flow Cytometry Core and by Leidos Biomedical Research Inc., especially the Laboratory Animal Sciences Program, Small Animal Imaging Program (Joe Kalen, Nimit Patel), Pathology Histotechnology Laboratory (Donna Butcher, Brad Gouker, and Elijah F. Edmondson), and Scientific Publications, Graphics and Media (Allen Kane). We thank Dr. Michael T. Lewis (BCM) for providing the PDX models and advice, and MaryBeth Hilton and Dr. Brad St.Croix for kind assistance with NSG mice. We thank Linda Miller and Suzanne Specht (NCI/LCDS) and Tiffany Dorsey (NCI, LHC) for outstanding assistance; Duncan Donohue and Tyler Malys (DMS, Inc.) for statistical analysis; Drs. Stan Lipkowitz, Deborah K. Morrison, and Namratha Sheshadri for suggestions during the study and comments on the manuscript; Drs. Wendy A. Woodward (MDACC), Mary L. Alpaugh (Rowan University), and D.K.M. for reagents (see also Methods); Drs. Xiaoping (Maggie) Wang, W.A.W, and M.L.A. for critical reading of the manuscript.

This research was supported by the Intramural Research Program of the NIH, National Cancer Institute, in part with Federal Funds under contract no. HHSN261200800001E and a pilot grant from METAvivor Research and Support, Inc., to K.B.. The Morgan Welch Inflammatory Breast Cancer Clinic is supported by the State of Texas Rare and Aggressive Breast Cancer Program. VP received support from the Isaac and Lucille Hay Graduate Fellowship. AJE received support from the BC Research Foundation (BCRF-20-048), the METAvivor Founder’s Award, and the NCI (U01CA217846).

