## Supplemental Information for "Stabilization of E-cadherin adhesions by COX-2/GSK3β signaling is a targetable pathway in metastatic breast cancer"

**CONTENT:**

- **Table S1**
- **Figures S1-S6**

**Table S1. Primers used for qPCR analysis.**

| GENE | FORWARD | REVERSE |
|---|---|---|
| *CDH1* | 5’-AGCAGAACTAACACACGGGG-3’ | 5’-ATACCGGGGGACACTCATGA-3’ |
| *CDKN1A* | 5’-ACCATGTGGACCTGTCACTGT-3’ | 5’-TTAGGGCTTCCTCTTGGAGAA-3 |
| *CEBPD* | 5’-GCCATGTACGACGACGAGA-3’ | 5’-TTGCTGTTGAAGAGGTCGG-3’ |
| *CTNNA1* | 5’-GGCAGCCAAAAGACAACAGG-3’ | 5’-TTACGTCCAGCATTGCCCAT-3’ |
| *CTNNB1* | 5’-TTGAAGGTTGTACCGGAGCC-3’ | 5’-GCCACCCATCTCATGTTCCA-3’ |
| *CTNND1* | 5'-TTGAGTGGGAATCGGTGCTC-3' | 5'-AGGAGGTCAGCTATGGCAGA-3' |
| *GAPDH* | 5’-AAGGTCGGAGTCAACGGATTTG-3’ | 5’-CCATGGGTGGAATCATATTGGAA-3’ |
| *IL6* | 5’-ACAAATTCGGTACATCCTC-3’ | 5’-GCAGAATGAGATGAGTTGT-3’ |
| *PTGS2* | 5’-GCTGTGGGGCAGGAAGTC-3’ | 5’-TTGGAATAGTTGCTCATCACC-3’ |
| *RPLP0* | 5’-GCAATGTTGCCAGTGTCTGTC-3’ | 5’-GCCTTGACCTTTTCAGCAAGT-3’ |

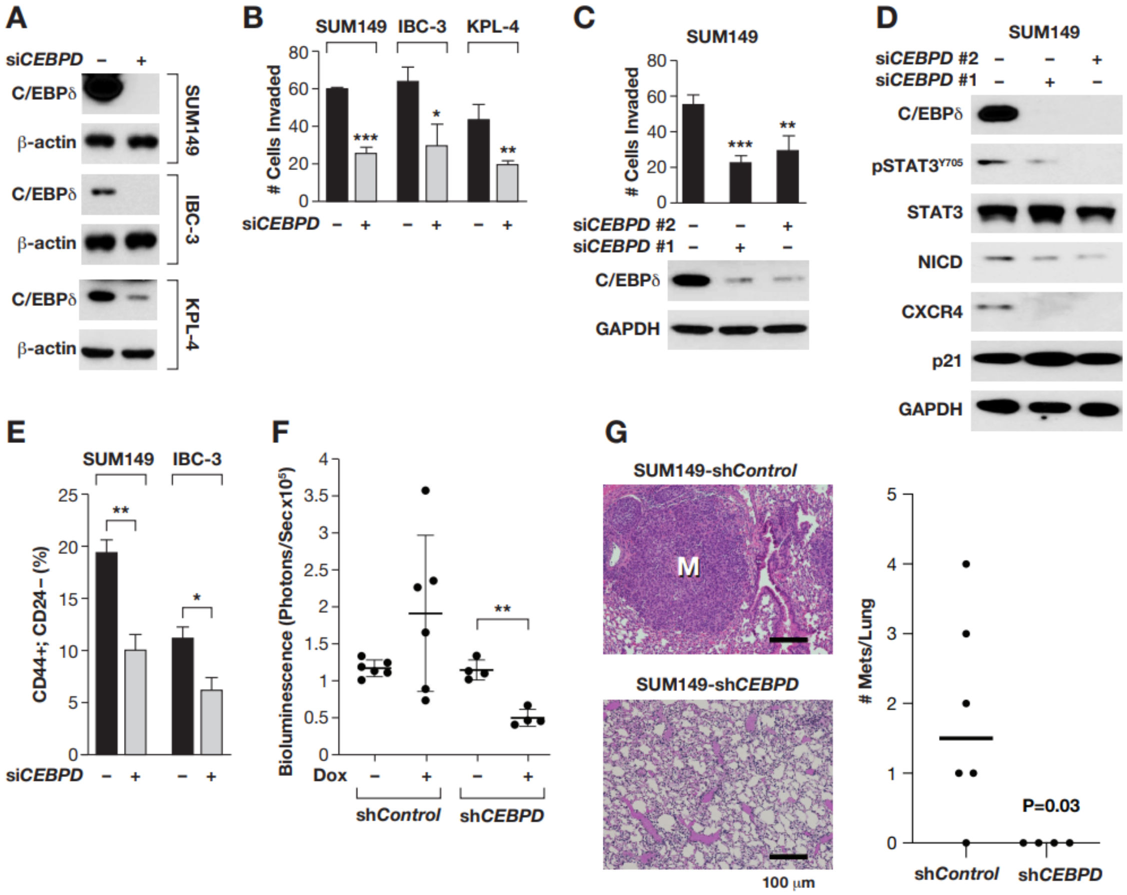


**Figure S1. C/EBPδ promotes malignant phenotypes in IBC cells**

**A**) Western analysis of C/EBPδ expression in SUM149, IBC-3 and KPL-4 cells 72 h after transfection with si*Control* (-) or si*CEBPD* oligos (+). **B**) Quantification of the number of cells invaded through transwell Matrigel by SUM149, IBC-3 and KPL-4 cells transfected with si*Control* (-) or si*CEBPD* (+) oligos (n=3, mean±SEM; **P*<0.05, ** *P*<0.01, ****P*<0.001 compared to si*Control*). **C**) Quantification of the number of cells invaded through transwell Matrigel by SUM149 cells transfected with si*Control* (-) or two independent si*CEBPD* (+) oligos (n=3, mean ±SEM; ***P*<0.01****P*<0.001 compared to si*Control*). Western analysis of C/EBPδ expression is shown below. **D**) Western analysis of the indicated proteins from SUM149 cells transfected with control (-) or two independent si*CEBPD* (+) oligos. **E**) Flow cytometric quantification of cells with CD44+:CD24– cell surface markers among SUM149 and IBC-3 cells transfected with control (-) or si*CEBPD* (+) oligos followed by culture in 2D for 3 days (n=3; mean ±SEM , * *P* <0.05, ** *P*<0.01). **F**) Quantification of bioluminescence (total Flux) in the lungs of mice with experimental metastases from SUM149 cells with doxycycline (Dox)-inducible shRNAs before (-) and 4 weeks after (+) treatment with Dox (***P*<0.01 by two-sided paired t-test, n=4-6). **G**) Light microscope images (left) of Hematoxylin and Eosin stained sections of lungs from mice as in panel F (M, metastasis) and quantification (right) of the number of tumor cell colonies (mets), n=4-6, P=0.03, two-sided unpaired Wilcoxon test.

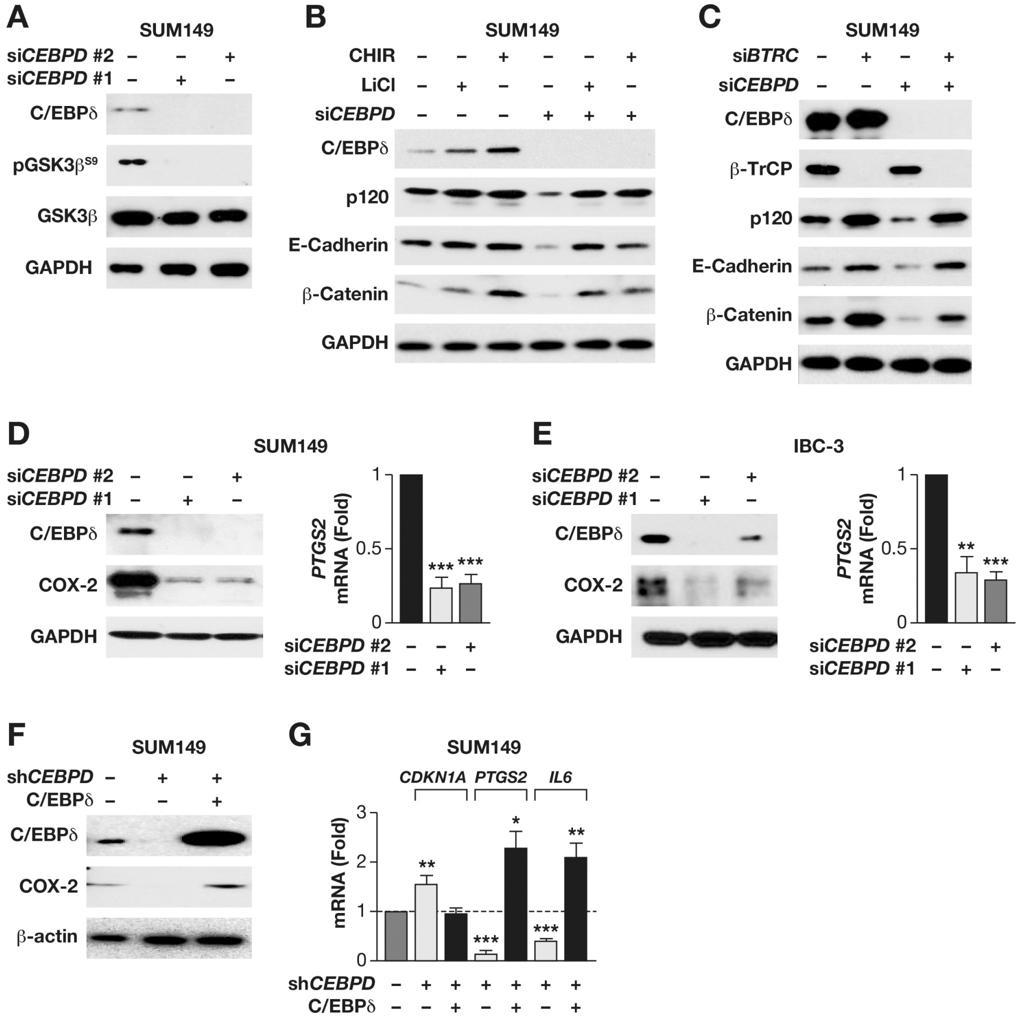


**Figure S2. C/EBPδ promotes COX-2 transcription.**
**A**) Western analysis of C/EBPδ, pGSK3β$^{S9}$, and GSK3β in emboli from SUM149 cells that were transfected with control or two independent si*CEBPD* oligos. **B**) Western analysis of the indicated proteins in emboli from SUM149 cells transfected with control (-) or si*CEBPD* oligos and treated with LiCl (10 mM) or CHIR (5 μM) for 6 h. **C**) Western analysis of the indicated proteins in emboli from SUM149 cells transfected with control (-) or si*CEBPD* along with si*BTRC* (β-TrCP) oligos. **D-E**) Western (left) and mRNA analyses by qPCR (right) of COX-2 (*PTGS2*) expression in (D) SUM149 and (E) IBC-3 cells 72 h after transfection with si*Control* or two independent si*CEBPD* oligos. n=3, mean±SEM, ** $P$<0.01, *** $P$<0.001 compared to si*Control*. **F**) Western analysis of COX-2 in SUM149 cells with stable expression of sh*Control* (-) or sh*CEBPD* (+) shRNA 72 h after transfection with vector (-) or C/EBPδ expression plasmid (+). **G**) qPCR analysis of the indicated mRNA's in cells as in panel F (n=3, mean±SEM; *$P$<0.05, ** $P$<0.01, *** $P$<0.001 compared to transfected with empty vectors).

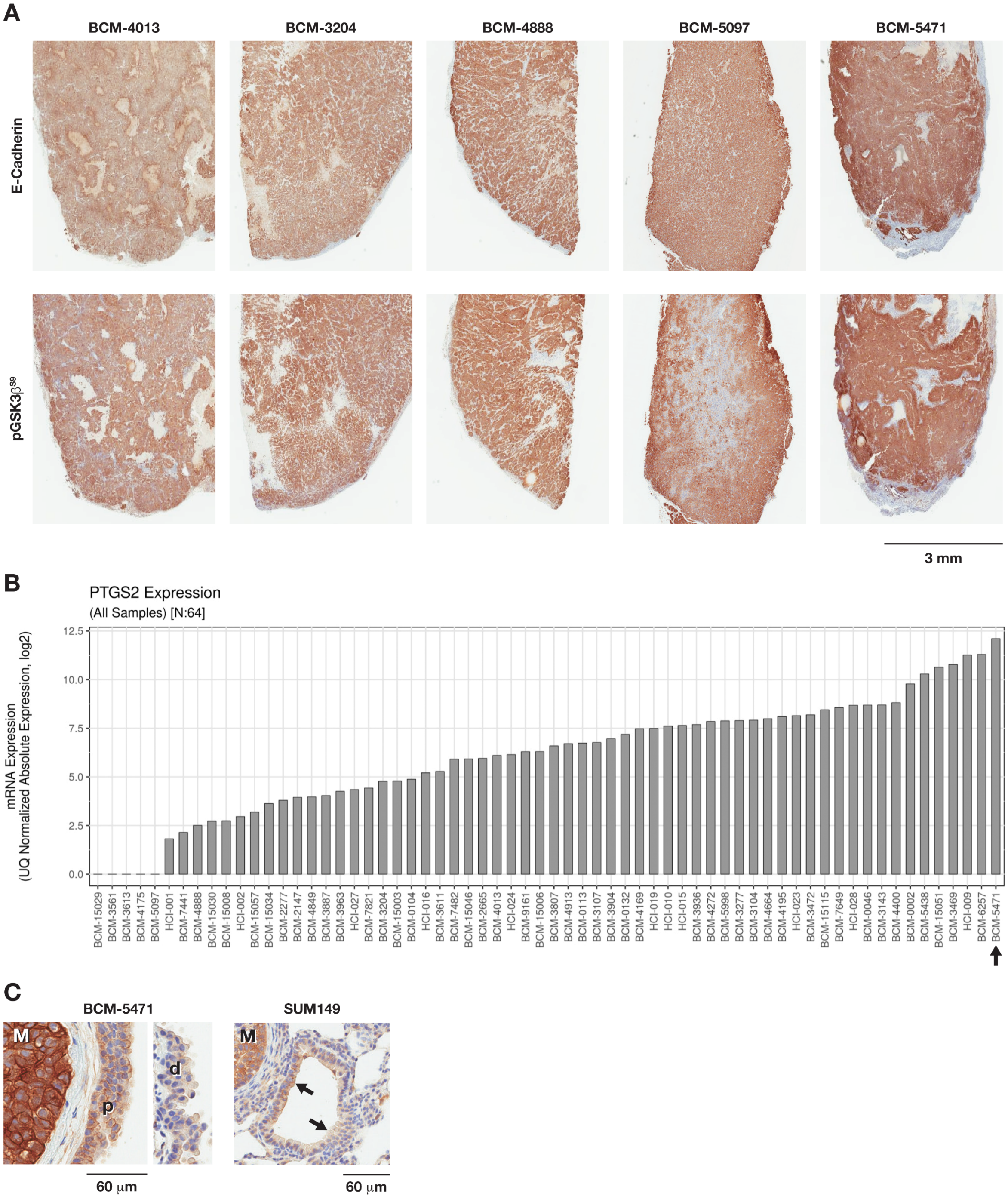


**Figure S3. E-Cadherin, COX-2 and/or pGSK3β$^{S9}$ expression analysis in breast cancer PDX primary tumors and lung metastases.**

**A**) Immunostaining of E-Cadherin and pGSK3β$^{S9}$ on serial sections of primary tumors from the indicated PDX models, representing TNBC (BCM-3204, -4013, 5471), ER+/HER2+ (BCM-4888), and ER+ (BCM-5097) subtypes. **B**) *PTGS2* mRNA data of BC PDX models as reported by the Patient-Derived Xenograft and Advanced In Vivo Models (PDX-AIM) Core at Baylor College of Medicine (https://pdxportal.research.bcm.edu). **C**) BCM-5471 and SUM149 lung metastases (M) with immunostaining of pGSK3β$^{S9}$ (p/d, proximal/distal, and black arrows indicate bronchial epithelium).

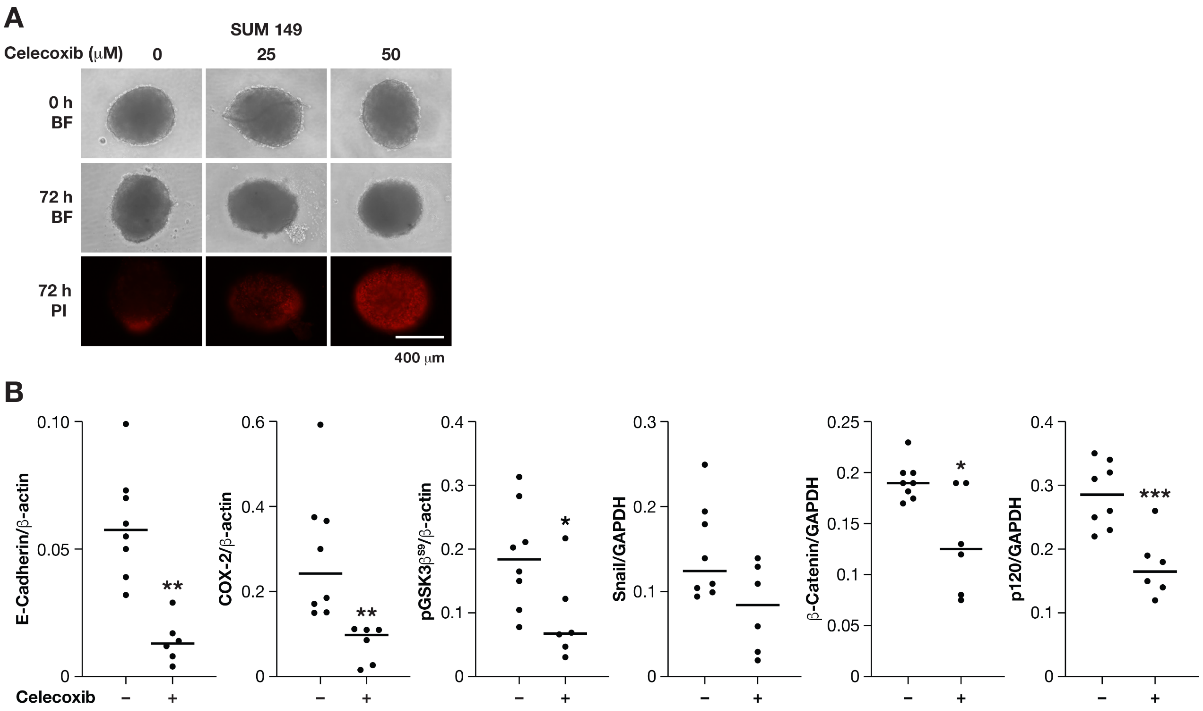


**Figure S4. Effect of Celecoxib on established emboli in vitro and tumors in vivo.**
**A**) Images of representative SUM149 cell emboli after 3 days of culture (0 h) and following another 72 h of treatment with celecoxib as indicated and stained with PI (BF, bright field; scale bar = 400 μm). **B**) Quantification of Western analysis shown in Figure 4I of BCM-5471 PDX tumors treated ± Celecoxib (n=6-8, mean±SEM; **P*<0.05, ** *P*<0.01 by unpaired two-sided Wilcoxon rank-sum test).

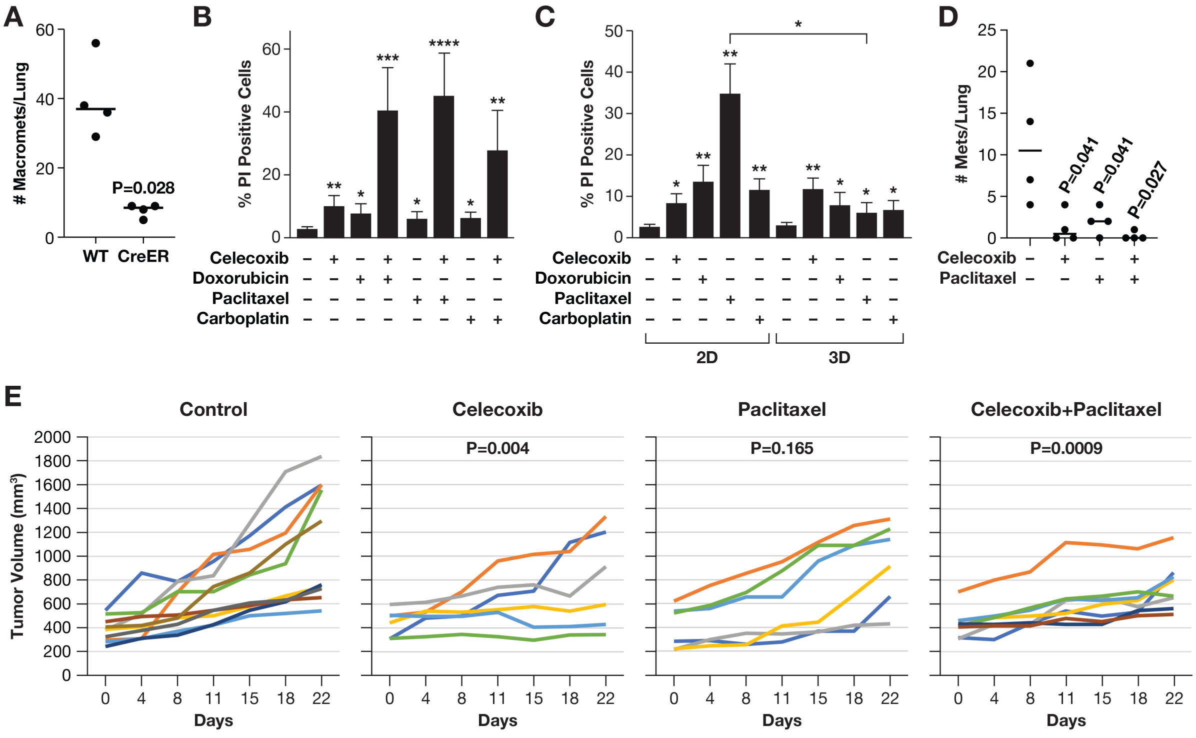


**Figure S5. Assessment of Celecoxib and paclitaxel drug sensitivity in vitro and in vivo.**
**A)** Quantification of macrometastases in lungs of mice injected with E-cadfl/fl or E-cadfl/fl; CreER expressing cells, followed 1 week later by tamoxifen injection to delete the E-cadherin gene, and harvested 3 weeks thereafter (n=4, P=0.0286 by two-tailed non-parametric t-test, Mann Whitney). **B**) Quantification of propidium iodide (PI) positive cells in emboli treated after 3 days of 3D culture for additional 3 days with individual drugs (50 µM celecoxib, 500 nM doxorubicin, 50 nM paclitaxel, 10 µM carboplatin) and/or combinations as indicated (n=3, mean±S.E.M.; *p<0.05, **p<0.01, ***p<0.001, ****p<0.0001 compared to DMSO control). **C**) Quantification of PI positive cells from cells cultured in 2D or 3D and treated as in panel B (n=3, mean±S.E.M.; *p<0.05, **p<0.01 compared to DMSO control or as indicated by bracket). **D**) Quantification of tumor cell colonies (mets) in lungs of mice injected with SUM149 cells and treated with celecoxib (1000 mg/kg chow) and/or paclitaxel (10 mg/kg i.v.), n=4, P as indicated by two-sided unpaired Wilcoxon test compared to untreated. **E**) Tumor volume measurements of BCM-5471 PDX's in mice treated with celecoxib (1000 mg/kg chow) and/or paclitaxel (10 mg/kg i.v.), n=6-10, P values were determined by a linear model of treatment by day interaction relative to control baseline.

**NM_005195.3 CEBPD mRNA**

```
AGGTGACAGCCTCGCTTGGACGCAGAGCCCGGCCCGACGCCGCCATGAGCGCCGCGCTCTTCAGCCTGGA   70
CGGCCCGGCGCGCGGCGCGCCCTGGCCTGCGGAGCCTGCGCCCTTCTACGAACCGGGCCGGGCGGGCAAG   140
CCGGGCCGCGGGGCCGAGCCAGGGGCCCTAGGCGAGCCAGGCGCCGCCGCCCCCGCCATGTACGACGACG   210
AGAGCGCCATCGACTTCAGCGCCTACATCGACTCCATGGCCGCCGTGCCCACCCTGGAGCTGTGCCACGA   280
CGAGCTCTTCGCCGACCTCTTCAACAGCAATCACAAGGCGGGCGGCGCGGGGCCCCTGGAGCTTCTTCCC   350
        TCGCCGACCTCTTCAACAG=siRNA#1

GGCGGCCCCGCGCGCCCCTTGGGCCCGGGCCCTGCCGCTCCCCGCCTGCTCAAGCGCGAGCCCGACTGGG   420
GCGACGGCGACGCGCCCGGCTCGCTGTTGCCCGCGCAGGTGGCCGCGTGCGCACAGACCGTGGTGAGCTT   490
GGCGGCCGCAGGGCAGCCCACCCCGCCCACGTCGCCGGAGCCGCCGCGCAGCAGCCCCAGGCAGACCCCC   560
GCGCCCGGCCCCGCCCGGGAGAAGAGCGCCGGCAAGAGGGGCCCGGACCGCGGCAGCCCCGAGTACCGGC   630
AGCGGCGCGAGCGCAACAACATCGCCGTGCGCAAGAGCCGCGACAAGGCCAAGCGGCGCAACCAGGAGAT   700
                                               pDEST_SIGMA=AACCAGGAGAT

GCAGCAGAAGTTGGTGGAGCTGTCGGCTGAGAACGAGAAGCTGCACCAGCGCGTGGAGCAGCTCACGCGG   770
GCAGCAGAAGT
GACCTGGCCGGCCTCCGGCAGTTCTTCAAGCAGCTGCCCAGCCCGCCCTTCCTGCCGGCCGCCGGGACAG   840
CAGACTGCCGGTAACGCGCGGCCGGGGCGGGAGAGACTCAGCAACGACCCATACCTCAGACCCGACGGCC   910
CGGAGCGGAGCGCGCCCTGCCCTGGCGCAGCCAGAGCCGCCGGGTGCCCGCTGCAGTTTCTTGGGACATA   980
GGAGCGCAAAGAAGCTACAGCCTGGACTTACCACCACTAAACTGCGAGAGAAGCTAAACGTGTTTATTTT   1050
                         siRNA#2=CCACTAAACTGCGAGAGAA
                           (Dox)SMARTchoice=CGAGAGAAGCTAAACGTGTTTATTTT

CCCTTAAATTATTTTTGTAATGGTAGCTTTTTCTACATCTTACTCCTGTTGATGCAGCTAAGGTACATTT   1120
GTAAAAAGAAAAAAAACCAGACTTTTCAGACAAACCCTTTGTATTGTAGATAAGAGGAAAAGACTGAGCA   1190
TGCTCACTTTTTTATATTAATTTTTACAGTATTTGTAAGAATAAAGCAGCATTTGAAATCGAAAAAAAAA   1260
```

**Figure S6.** ***CEBPD* targeting sequences.**
The human *CEBPD* sequence (NM_005195.3) is shown along with the aligned sequences of each siRNA and shRNA (pDEST_SIGMA) used in this study. The translation start and stop codons are indicated by shaded boxes.
